# MicroRNA variants and HLA-miRNA interactions are novel rheumatoid arthritis susceptibility factors

**DOI:** 10.1101/2020.08.25.264515

**Authors:** Shicheng Guo, Yehua Jin, Jieru Zhou, Qi Zhu, Ting Jiang, Yanqin Bian, Runrun Zhang, Cen Chang, Lingxia Xu, Jie Shen, Xinchun Zheng, Yi Shen, Yingying Qin, Jihong Chen, Xiaorong Tang, Peng Cheng, Qin Ding, Yuanyuan Zhang, Jia Liu, Qingqing Cheng, Mengru Guo, Zhaoyi Liu, Weifang Qiu, Yi Qian, Yang Sun, Yu Shen, Hong Nie, Steven J Schrodi, Dongyi He

## Abstract

**Objective:** Althogh Genome-wide association studies have identified >100 variants for rheumatoid arthritis (RA),the reported genetic variants only explain <40% of RA heritability. We conducted a systemic association study between common East-Asian miRNA SNPs with RA in a large Han Chinese cohort to explain missing heritability and identify miRNA epistatic interactions.

**Methods:** 4 HLA SNPs (HLA-DRB1, HLA-DRB9, HLA-DQB1 and TNFAIP3) and 225 common SNPs located in miRNA which might influence the miRNA target binding or pre-miRNA stability were genotyped in 1,607 rheumatoid arthritis and 1,580 matched normal individuals. A meta-analysis with previous GWAS studies (4,873 RA cases and 17,642 controls) was performed to discovery another novel miRNA RA-associated SNPs.

**Results:** 2 novel SNPs including rs1414273 (miR-548ac, OR=0.84, P=8.26×10-4) and rs2620381 (miR-627, OR=0.77, P=2.55×10-3) conferred significant association with RA. Individuals carried 8 risk alleles showed 15.38 (95%CI: 4.69-50.49, P<1.0×10-6) times more risk to be affected by RA. In addition, rs5997893 (miR-3928) showed significant epistasis effect with rs4947332 (HLA-DRB1, OR=4.23, P=0.04) and rs2967897 (miR-5695) with rs7752903 (TNFAIP3, OR=4.43, P=0.03). Finally, we demonstrated targets of the significant miRNAs showed enrichment in immune related genes (P=2.0×10-5) and FDA approved drug target genes (P=0.014).

**Conclusions:** 6 novel miRNA SNPs including rs1414273 (miR-548ac, P=8.26×10-4), rs2620381 (miR-627, P=2.55×10-3), rs4285314 (miR-3135b, P=1.10×10-13), rs28477407 (miR-4308, P=3.44×10-5), rs5997893 (miR-3928, P=5.9×10-3) and rs45596840 (miR-4482, P=6.6×10-3) were confirmed to be significantly associated with RA in a Chinese population. Our study suggests that miRNAs might be interesting targets to accelerate the understanding of the pathogenesis and drug development for rheumatoid arthritis.

## Introduction

Rheumatoid arthritis (RA) is a chronic inflammatory disorder caused by the interaction between multiple factors including genetics, epigenetics and the environment.^1–3^ Twin studies estimate that an RA heritability of ~60%.^4^ ^5^ In the past 15 years, GWAS have identified several hundred RA-associated variants. However, the reported genetic variants only explain <40% of RA heritability.^5–8^ Furthermore, studies within non-European-derived samples will aid our understanding of RA molecular pathophysiology as different variants and, potentially, different disrupted pathways may be present in those sample sets. Additionally, studying SNPs in specific functional classes of genes or motifs may reveal specific pathogenic mechanisms not previously discovered through GWAS.^9^ GWAS results indicate that 90% of disease-associated variants are located in non-coding regions, indicating regulatory elements may play important roles in complex disease etiology, including RA.^10^ Across regulatory elements, microRNAs are important targets to interrogate as miRNA SNPs may modify expression at numerous genes. Moreover, miRNAs play pivotal roles in both innate and adaptive immunity,^11^ ^12^ sex-specific effects,^13^ and disease onset.^14^ ^15^

Recently, several preliminary RA-association studies focused on miRNAs have been conducted in Asian,^16–19^ European^20^ and African^21^ samples. However, these studies have very limited sample sizes and only included a subset of miRNAs without genotyping all miRNA common SNPs. Additionally, the previous RA GWAS in Han Chinese was conducted in 2014^22^ using an array with poor coverage of miRNA SNPs. Therefore, we conducted an exhaustive study of functional miRNA SNPs in RA. miRbase annotates 1,920 primary miRNA and 2,883 mature miRNAs. From dbSNP153, there are >45,705 SNPs located in primary miRNAs and 17,570 in mature miRNA;^23^ however, only 733 of these SNPs segregate alleles at moderate-high frequency (i.e. are common SNPs with MAF>1%) within primary miRNAs regions. Within specific populations, this number is expected to be substantially reduced to ~200 SNPs, providing an opportunity to use a multiplex genotyping assay in our RA samples. Hence, we conducted a systemic association study between common East-Asian miRNA SNPs with RA in a large Han Chinese cohort to identify novel miRNA SNPs and miRNA epistatic interactions.

## Methods

### Precision Medicine Research Cohort in Shanghai Guanghua Hospital

Guanghua Hospital Precision Medicine Research Cohort (PMRC) is a hospital-based longitudinal cohort to investigate risk factors, genetic susceptibility, pharmacogenetics for rheumatology diseases such as RA, osteoarthritis and ankylosing spondylitis. Healthy individuals are derived from those with an annual physical exam without rheumatological disease. Currently, PMRC has enrolled >30,000 disease patients and 10,000 healthy individuals as controls. We randomly selected 3,223 individuals including 1,625 seropositive (RF+ and anti-CCP+) RA and 1,598 controls from the PMRC. Written consent was collected prior to enrollment. Non-Han Chinese individuals were excluded from the study to avoid confounding by population stratification. All cases fulfilled the 2010 European League against Rheumatism–American College of Rheumatology criteria or 1987 American College of Rheumatology revised criteria for RA. All healthy controls were required do not have personal or family history of ankylosing spondylitis, rheumatoid arthritis, osteoarthritis, type 1 and 2 diabetes, chronic infection or common cancers. We did not find significant case/control difference between gender, age, smoking, BMI after correction for multiple testing. This study was reviewed and approved by the Institutional Review Board of Guanghua Hospital (No: IRB12018-K-12) and all methods were performed in accordance with the relevant guidelines and regulations. The demographic and clinical characteristics of the whole samples are presented in **Table S1**.

### DNA extraction, SNP selection and Genotyping

Genomic DNA was extracted from peripheral blood using the CB-kit (CoWin Biosciences, CWBIO, China) according following manufacturer’s instructions. Allelic discrimination was automated using the manufacturer’s software which has been widely-described in previous studies.^24^ Internal positive and negative control samples were used and a test of Hardy-Weinberg equilibrium was employed to assess the genotyping quality (SNPs exceeding HWE p-value cutoff were excluded from RA association testing). Cases and controls were required to have a family history of at least three generations of residency in Shanghai or neighboring regions. The SNaPshot Multiplex System (Applied Biosystem,USA) was used for the genotyping which has been widely used in our previous studies.^25^ ^26^

To select SNPs for subsequent association testing, human microRNAs were downloaded from miRBase methjods (Release 22.1). Mapping of SNPs within dbSNP (dbSNP153, 08/08/2019) was performed for human microRNA genomic regions, resulting in 40,602 SNPs. To select common SNPs (MAF>0.01), we collected the allele frequency for all the SNPs from Gnomad,^27^ Asian100K^28^ and the 1000 Genomes dataset.^10^ The minor allele frequency was required to be higher than 0.01 in all datasets. [t]^29^ Only biallelic SNPs were included in the analysis, while tri-allelic SNPs were excluded from this study. Following filtering, 243 SNPs were obtained. In order to decrease the genotyping cost, one SNP from pairs of SNPs with high LD (R^2^>0.8) were removed and only one of them was included in the genotyping assay. A total of 233 SNPs were genotyped. GRCH37/hg19 was used to determine genomic positions and R^2^>0.6 as the cut-off to selected imputed SNPs of high quality and located in miRNA regions. The GAsP reference panel showed higher imputation accuracy (93%-95%) compared with 1000 Genome Asian panel (<90%).^30^ Four SNPs within known GWAS associations with RA were included: rs9268839 (*HLA-DRB9*),^31^ ^32^ rs9275376 (*HLA-DQB1*), ^33–36^ rs4947332 (tag-SNP for *HLA-DRB1*04:07*) and rs7752903 (*TNFAIP3*).^31^ ^32^ In addition, a panel of five ancestry informative markers (negative control) were selected to estimate the potential effects of population stratification: rs174583 (11q12.2), rs11745587 (5q31.1), rs521188 (1p31.3) and rs7740161 (6q23.2). These SNPs were derived from a previous Han Chinese population study.^37^ These SNPs are informative for interrogating population structure between South and North Han Chinese since the Shanghai region is a mixed population from different regions in China (**Figure 1A**).

**Figure 1.**
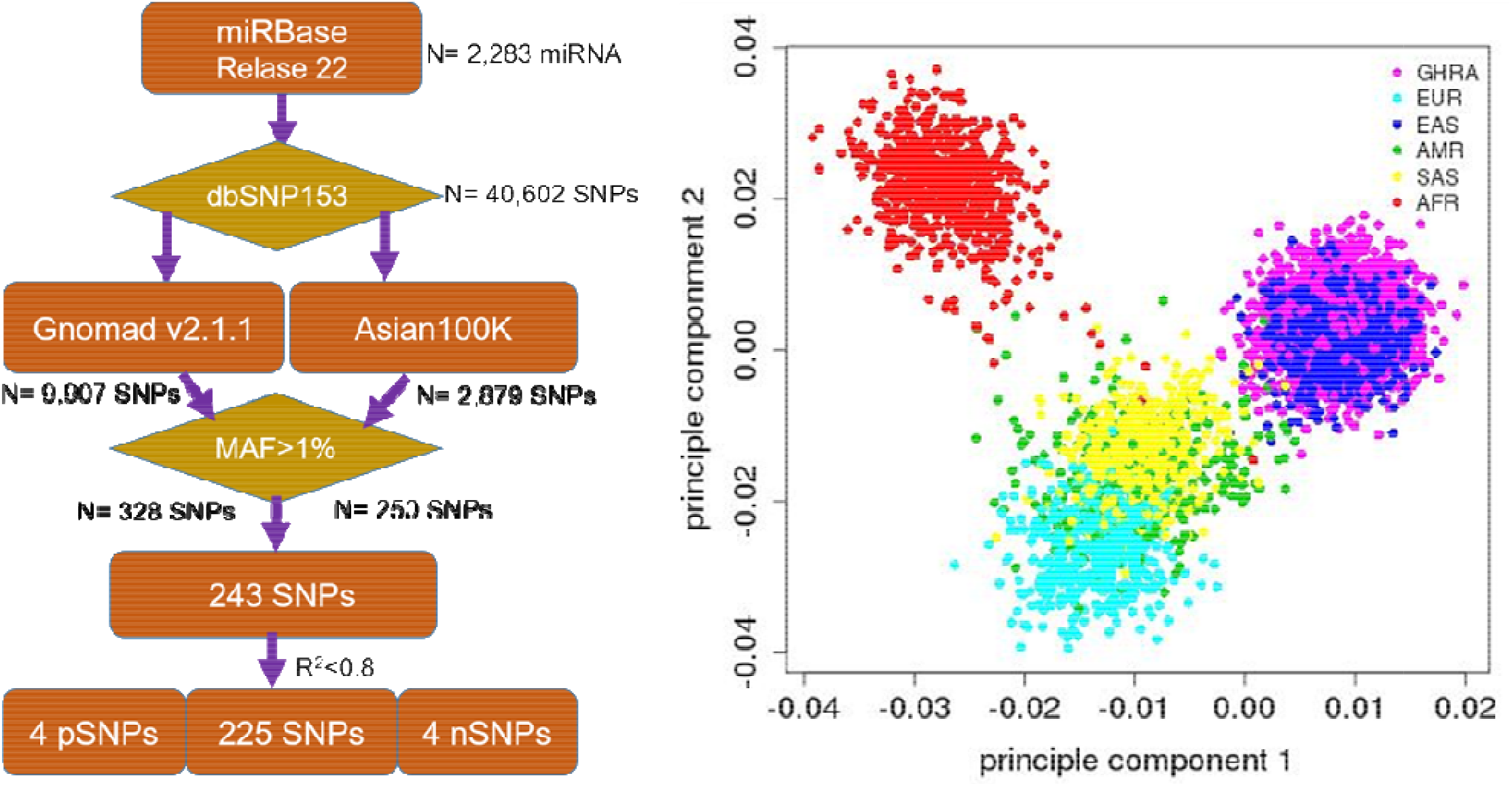
Flowchart for SNP and sample selection as well as population ancestry of GHRA cohort. (A) human microRNAs were downloaded from miRBase (Release 22.1). We mapped all the dbSNP153 SNPs into human Asian 100K data with MAF>1%. 4 positive control (*HLA-DRB1, HLA-DRB9, HLA-DQB1* and *TNFAIP3*) and 4 ancestry markers (South and North Han Chinese) were added for the quality control. (B) PCA analysis to show the population ancestry of Guanghua Rheumatoid Arthritis cohort (GHRA) with 1000 Genome dataset. Symbols are as same as Super Population Code (SPC) from 1000 genome project. AFR, African; AMR, Ad Mixed American; EAS, East Asian; EUR, European; SAS, South Asian.

### Quality Control and Power analysis

For genotyping QC, ~2.5% samples (40 RA and 32 normal) were randomly selected for repeat genotyping. The concordance between initial and replicate genotyping was 99.98%. SNP imputation was accomplished through the Michigan Imputation Server (https://imputationserver.sph.umich.edu/index.html) with East Asian samples from the Genome Asian Pilot (GAsP). Missing ratio between case and control differential test and HWE in control samples were evaluated. SNPs were removed when P<0.01.

In the power analysis, given the prevalence of RA (p=0.01), case (n=1,607) and control (n=1,580) size, risk allele frequency (f=0.1) and genetic effect size (OR=1.5), the power for multiplicative model, additive model, dominant models were 0.856, 0.823 and 0.742, respectively using a significance level of alpha=5×10^−03^. In addition, with our design and the above effect size, we calculate that SNPs with MAF of 0.025 can attain a statistical power of 80% with nominal p<0.05. Therefore, our study could provide reliable genetic association results for the majority of the alleles we have included given that the MAF of 201 (86.3%) SNPs are >10% and MAF of 96% included SNPs exceed 2.5%.

### Bioinformatics analysis of miRNA regulatory networks and enrichment analysis

To understand regulatory networks driven by miRNAs and the overlap with RA GWAS genes and immune system genes, a miRNA regulatory network analysis was performed with miRDB.^38^ Genes predicted to be regulated by significant miRNAs were calculated among 123 previously-identified RA GWAS genes, 4,723 immune system genes collected by InnateDB database^39^ and 373 FDA-targeted pharmacogenes. A target sore threshold was set to 95 in miRDB to identify reliable gene targets for the miRNAs. Cytoscape was employed to construct and visualize the miRNA-mRNA network. We calculated a hypergeometric test of the enrichment in miRNA targets within RA GWAS significant genes, immune-related genes, and FDA approved drug targets. A random sampling technique was applied to assess the null distribution of p-values (n=50,000 iterations). 37,875 total genes in the genome (GENCODE v32) were assumed for the calculations. Enrichment was measured with fold-change (FC) and p-values were calculated by summing the frequency of more extreme values in the null distribution of p-values.

### Association Analysis, Power Calculations, Epistasis Analysis and Cumulative Risk Analysis

RA Association for each SNP were calculated and OR estimated (95% CI) using Chi-square test, Fisher’s Exact test (N<5), Cochran-Armitage Trend test and Bayesian logistic regression (BLR)^40^ adjusted for gender, age, BMI, smoking and drinking history, using the bayesm (version: 3.1.4) package.^40^ To explore the power of the study, a Monte Carlo simulation was performed under different models including dominant, recessive and additive models, additive models, using a Chi-square test or Bayesian logistic regression model. SNP-SNP interactions were analyzed by using traditional point-wise interaction analysis based on SNPassoc,^41^ logistic test,^42^ “fast-epistasis” in Plink.^43^ We applied *Y ~ b0 + b1*A + b2*B + b3*AB + e* to detect epistasis effect between SNP A and SNP B when b3 not equal 0 (null hypothesis: b3 equal 0). A *P*<0.001 was considered significant. An RA risk prediction model was fitted with the 6 most significant SNPs (rs1414273, rs4947332, rs9268839, rs9275376, rs7752903 and rs2620381) by both Chi-square and logistic regression test to adjust by above-mentioned covariates. Q-value received after multiple test correction was conducted through a False Discover Rate (FDR) implemented in R with the function p.adjust).^44^ SNP derived miRNA target gain or loss were imputed by miRNA-SNP database^45^ with the default setting. All the statistical analyses were performed with R (version: 3.6.1).

## Results

### SNP Selection, Genotyping and Population Structure

Overall, the genotyping generated high quality data with 94% of the SNPs showing high genotyping quality. Seven of the SNPs were removed from being promoted to the RA association evaluation due to a genotyping ratio less than 99.0%. The missing ratio between case and control was calculated. Hardy-Weinberg equilibrium in control samples was evaluated and another 3 SNPs were removed when P<0.01. Finally, 223 SNPs in 1,607 rheumatoid arthritis and 1,580 normal individuals were used in the association analysis (genotyping rate=98.88%). PCA analysis based on these SNPs showed our samples clustered with the East Asian population (**Figure 1B**). Furthermore, we did not identify significant age, BMI, drinking and smoking history differences between cases and controls (P>0.01, **Table S1**). However, as a conservative measure, association testing of all the variants included age, gender, BMI, drinking and smoking as covariates in the Bayesian logistic regression (BLR)-based tests.

### Novel RA-associated SNPs located in *miRNA-548* and *miRNA-627*

Applying the Bayesian logistic regression model adjusted for covariates, we identified 6 significant SNPs located in *HLA-DRB9* (rs9268839, P= 3.95×10^−27^), *HLA-DRB1* (rs4947332, P=2.78×10^−4^), *HLA-DQB1* (rs9275376, P=2.65×10^−20^), *TNFAIP3* (rs7752903, P=2.33×10^−4^), *miR-548ac* (rs1414273, P=8.26×10^−4^) and *miR-627* (rs2620381, P=2.55×10^−3^) and 15 additional SNPs with P<0.05 (**Table 1**). As expected, all the SNPs located in *HLA-DRB1, HLA-DRB9, HLA-DQB1* and *TNFAIP3* were significantly associated with RA status, which is consistent with previous GWAS studies. We found all the population markers including rs174583 (FDR=0.88), rs11745587 (FDR=0.98), rs521188 (FDR=0.99) and rs7740161 (FDR=0.48) were not significant in the association test between RA and control. Additionally, two miRNA SNPs located in *miR-548ac* (FDR=0.01) and *miR-627* (FDR=0.045) were significantly associated with RA after FDR adjustment for multiple testing (P<0.05, FDR). We also conducted mode of inheritance-based association analyses (**Table 2**) and Cochran-Armitage Trend test-based association analyses (**Table S2**). We found a dominant model of two miRNA SNPs (rs1414273 and rs2620381) showed highly significant association between the alleles and RA status. To combine previous Han Chinese association data with this study, we also implemented a metaanalysis in which 4,873 RA cases and 17,642 normal individuals from the East Asian Population (EAP) were included. The meta-analysis resulted in an additional 4 significant SNPs (**Table 3, Figure 2**) including rs4285314 (miR-3135b, FDR=1.10×10^−13^), rs28477407 (*miR-4308,* FDR=3.44×10^−5^), rs5997893 (*miR-3928,* FDR=5.9×10^−3^) and rs45596840 (*miR-4482,* FDR=6.6×10^−3^). Finally, we also evaluated whether rs1414273 (*miR-548ac*) and rs2620381 (*miR-627*) are significant RA susceptibility variants independent of HLA variants. We found that the association at rs1414273 (OR=1.18, P=1.03×10^−3^) and rs2620381 (OR=1.31, P=2.55×10^−3^) remained significant following adjustment for HLA alleles.

**Figure 2.**
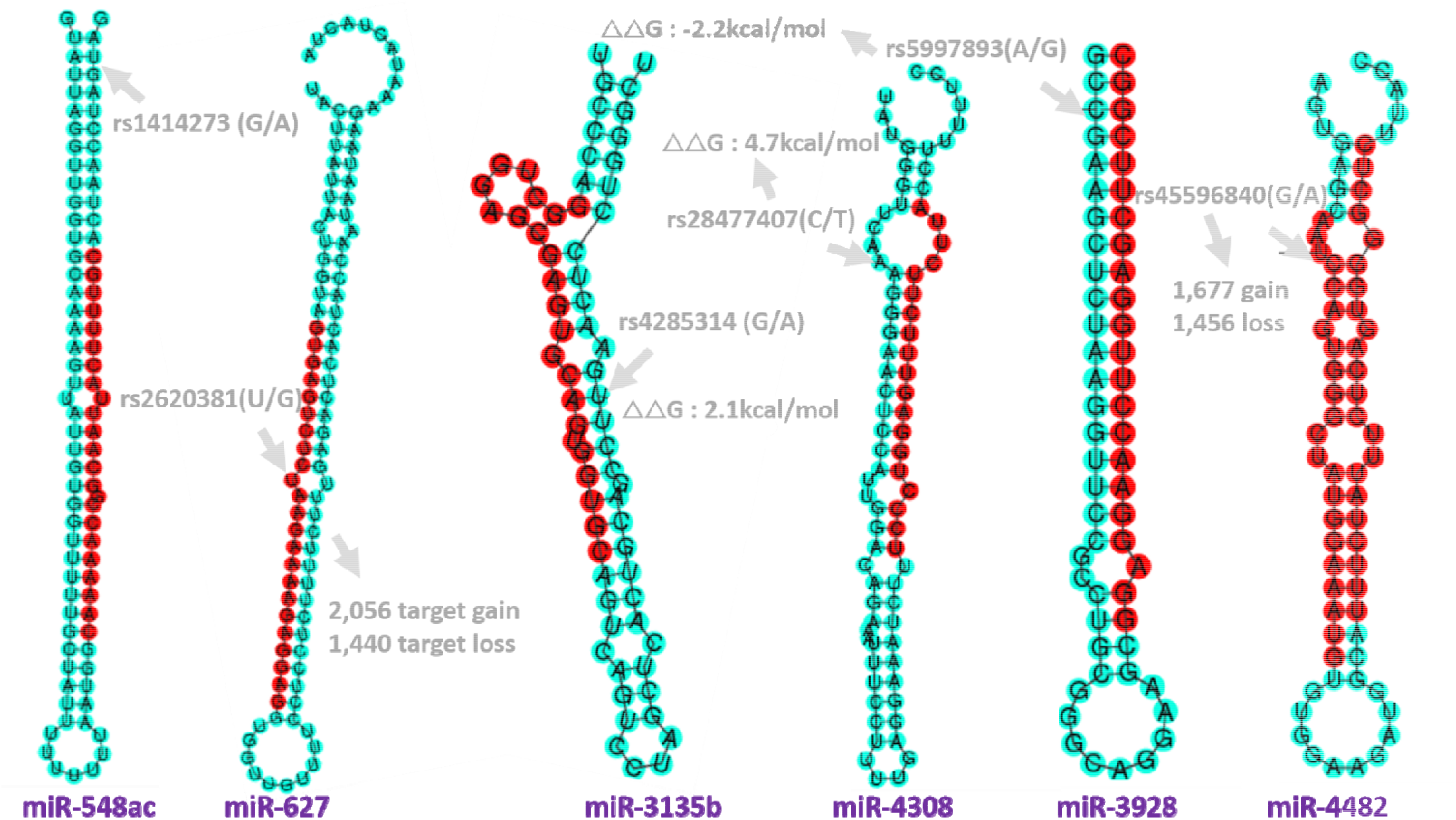
Genomic position and functional annotation to the significant SNPs in miRNAs. rs1414273 (C/T) in *miR-548ac*; rs2620381 (A/C) in *miR-627-5p* which cause 2,056 target gain and 1,440 target loss. rs4285314(G/A) in miR-3135b; rs28477407(C/T) in *miR-4308;* rs5997893 (A/G) in *miR-3928-5p* and rs45596840 (G/A) in *miR-4482-5p* which will cause 1,677 target gain and 1,456 target loss. Target gain or loss were imputed by miRNA-SNP database^45^ (see method).

**Table 1.**
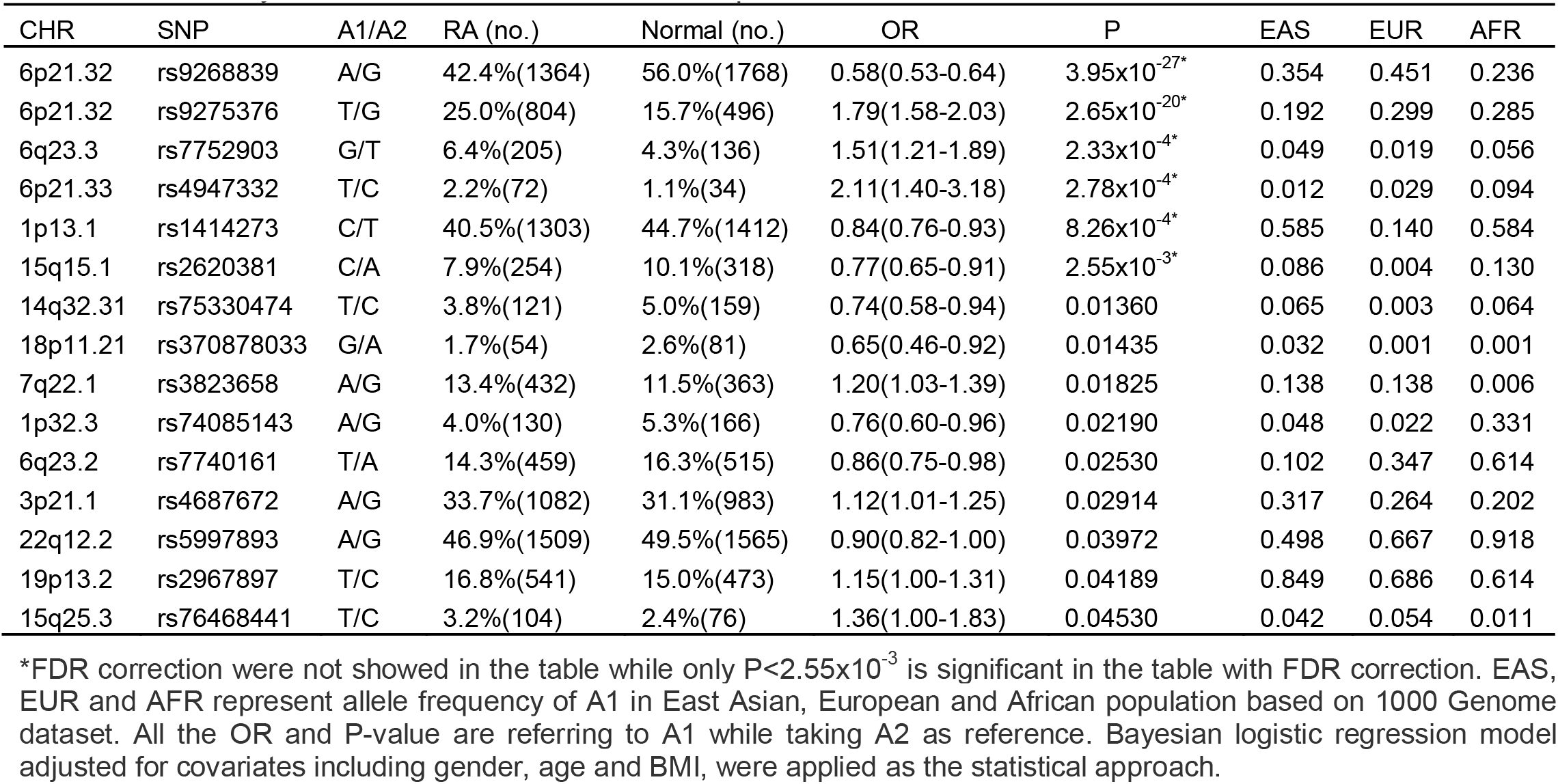
Summary of risk alleles, 1000 Genome frequencies and SNPs associated with rheumatoid arthritis. *FDR correction were not showed in the table while only P<2.55×10^−3^ is significant in the table with FDR correction. EAS, EUR and AFR represent allele frequency of A1 in East Asian, European and African population based on 1000 Genome dataset. All the OR and P-value are referring to A1 while taking A2 as reference. Bayesian logistic regression model adjusted for covariates including gender, age and BMI, were applied as the statistical approach.

**Table 2.**
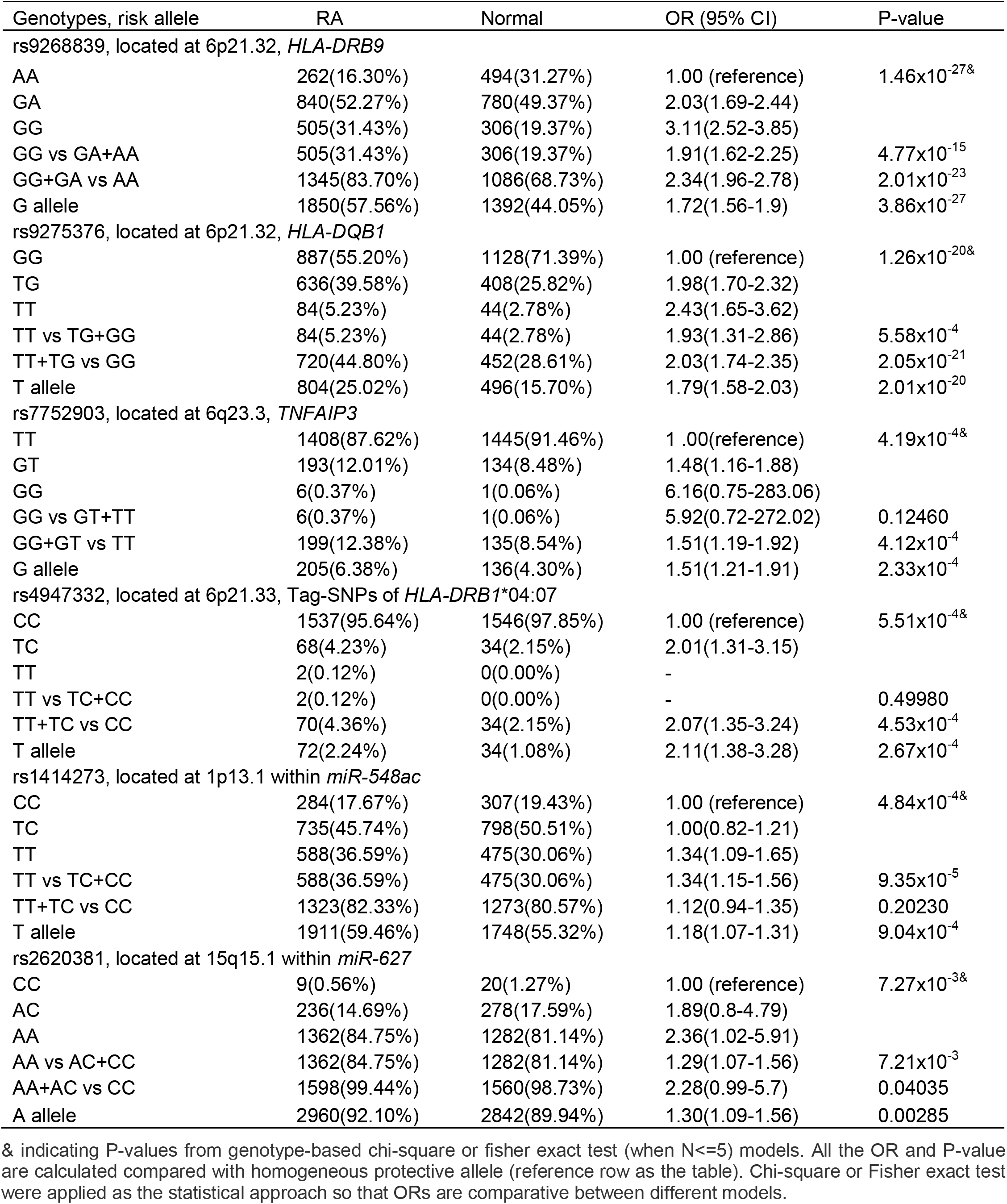
Association statistics for rheumatoid arthritis susceptibility and genetic variants. & indicating P-values from genotype-based chi-square or fisher exact test (when N<=5) models. All the OR and P-value are calculated compared with homogeneous protective allele (reference row as the table). Chi-square or Fisher exact test were applied as the statistical approach so that ORs are comparative between different models.

**Table 3.**
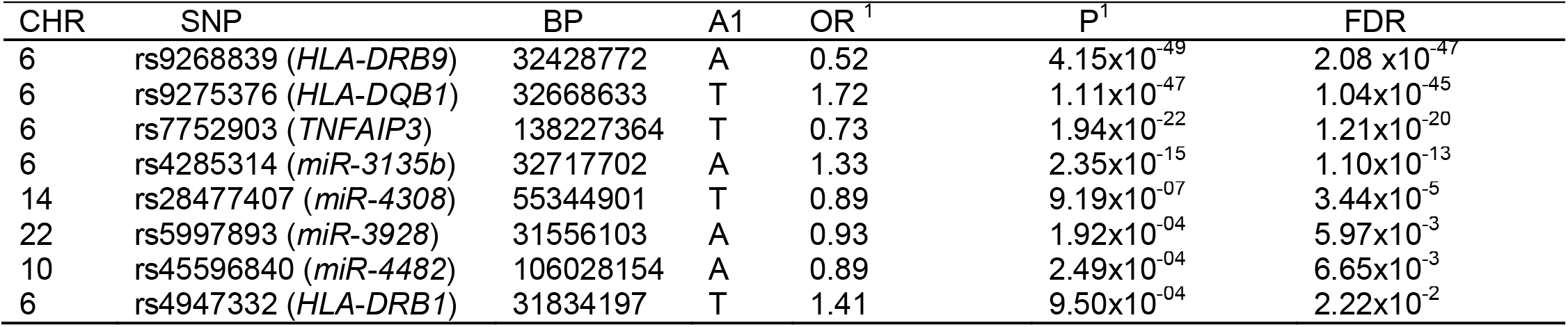
meta-analysis between our study and previous East-Asian population-based RA GWAS study.

### Epistasis analysis to identify miRNA interactions

miRNAs play multiple regulatory roles in both modification of target gene expression and larger regulatory networks. In order to identify epistatic effects between miRNAs and their role in the susceptibility of RA, we applied an epistasis analysis to reveal the interaction between above identified miRNA SNPs. We found 19 SNP-SNP epistatic interactions with P<7.3×10^−4^, indicating significant interactions (**Table S3**). We found 10 SNP-SNP pairs showed significant strengthened interaction (OR>1) while nine SNP-SNP pairs showed impaired interaction (OR<1). In addition, enhanced risk epistasis was also identified across HLA alleles (rs4947332 in *HLA-DRB1 and* rs9275376 in *HLA-DQB1*) between HLA and miRNAs (*HLA-DRB1* and miR-3928), and between *TNFAIP3* and miR-5695 (**Table 4**). A significant positive interaction between rs4947332 (*HLA-DRB1*) and rs5997893 (miR-3928) with a significantly inflated OR=2.83 (95%CI: 1.75-4.58, P=1.36×10^−5^, **Table 4**) for double risk allele carriers, indicating the importance of HLA and non-HLA genetic variation interaction in RA susceptibility.

**Table 4.** Epistasis analysis to identify SNP-SNP interaction in rheumatoid arthritis susceptibility. Five significant epistasis effect were selected to show the increased or decreased risk effect. rs4947332 located in C2 upstream (*HLA-DRB1);* rs9275376 located in *HLA-DQB1* upstream; rs5997893 located in *miR-3928;* rs7752903 located in downstream of TNFAIP3 and rs2967897 located in *miR-5695.*

| Epistasis | SNP Genotypes |  | RA | Control | OR | P |
| --- | --- | --- | --- | --- | --- | --- |
| rs4947332& rs9275376 | rs4947332 | rs9275376 |  |  |  |  |
|  | CC | GG | 885 | 1119 | 1.00 (reference) | - |
|  | CT+TT | GG | 2 | 9 | 0.38(0.10-1.38) | 0.164 |
|  | CC | GT+TT | 652 | 427 | 1.93(1.66-2.24) | 6.59x10<sup>-18</sup> |
|  | CT+TT | GT+TT | 68 | 25 | 3.35(2.12-5.31) | 6.70x10<sup>-8</sup> |
| rs4947332& rs5997893 | rs4947332 | rs5997893 |  |  |  |  |
|  | CC | GG | 330 | 392 | 1.00 (reference) | - |
|  | CT+TT | GG | 9 | 9 | 1.19(0.49-2.89) | 0.821 |
|  | CC | GT+TT | 1207 | 1154 | 1.24(1.05-1.47) | 0.012 |
|  | CT+TT | GT+TT | 61 | 25 | 2.83(1.75-4.58) | 1.36x10<sup>-5</sup> |
| rs7752903& rs2967897 | rs7752903 | rs2967897 |  |  |  |  |
|  | TT | CC | 953 | 1050 | 1.00 (reference) | - |
|  | TG+GG | CC | 148 | 93 | 1.75(1.33-2.29) | 5.76x10<sup>-5</sup> |
|  | TT | CT+TT | 455 | 395 | 1.27(1.08-1.49) | 0.00370 |
|  | TG+GG | CT+TT | 51 | 42 | 1.33(0.88-2.01) | 0.20705 |

### Cumulative analysis revealed increased risk effect on rheumatoid arthritis

To estimate the combined effect of the 6 risk alleles (rs1414273T, rs4947332T, rs9268839G, rs9275376T, rs7752903G and rs2620381A) on RA risk, the individual accumulation of risk alleles was treated as an ordinal variable in a logistic regression analysis adjusted by BMI, age and gender. As anticipated, we found the RA risk showed a positive correlation with cumulative number of risk alleles (OR=1.4, P = 2.0×10^−16^, Z=12.54, SE=0.027, **Table 5**). In our dataset, the largest subgroup of cases carries five risk alleles (25.6% of RA samples) while the largest subgroup of controls carries only 4 risk alleles (29.4% of control samples). The OR for RA status for carriers with eight risk alleles (2.9% of RA population) was 15.38-fold increased over individuals with only 1 risk alleles (10.76% of normal population).

**Table 5.**
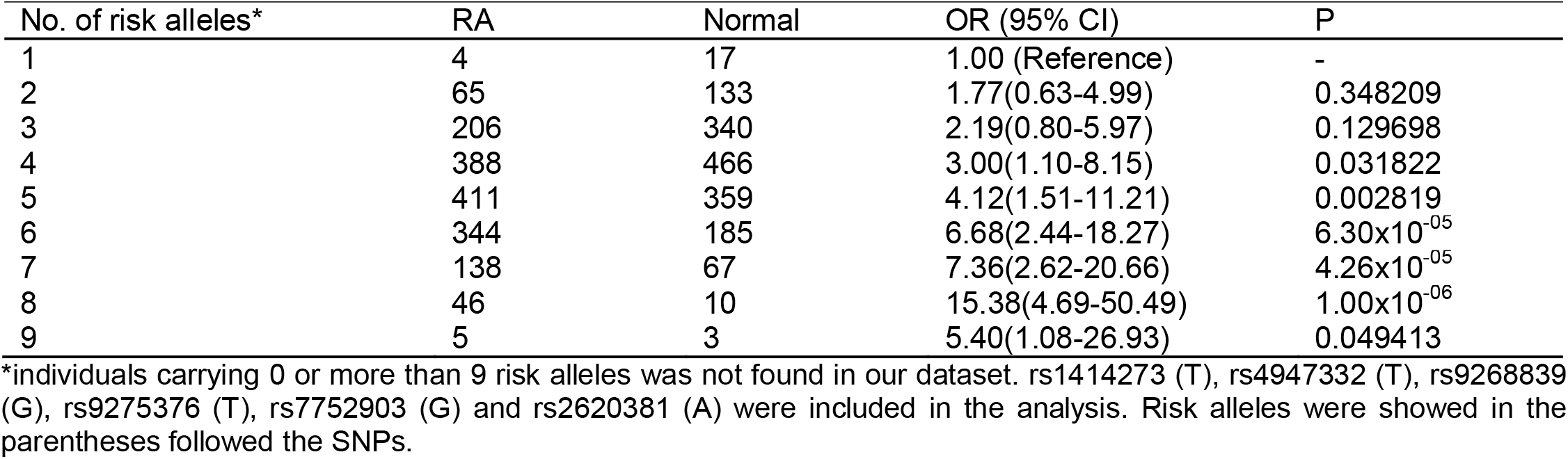
Cumulative risk effect of rheumatoid arthritis increased as more risk alleles carried by the patients. *individuals carrying more than 9 risk alleles were not found in our dataset. rs1414273 (T), rs4947332 (T), rs9268839 (G), rs9275376 (T), rs7752903 (G) and rs2620381 (A) were included in the analysis. Risk alleles were showed in the parentheses followed the SNPs.

### Regulatory network constructed by significant miRNAs and RA GWAS associated genes

To construct regulatory networks between miRNAs and genes, we predicted target genes for the significant miRNAs identified in our study. First, we predicted candidate genes of our significant miRNAs and then mapping these targets to overall immune related genes collected by InnateDB. We found target genes of RA-associated miRNAs were significant enriched in the immune related gene category (P<2.2×10^−16^, empirical P=2.0×10^−5^ and FC=1.43, **Figure 3A, Figure S1A**). In addition, we found *miR-548ac* has 12 target genes overlapping with previous GWAS-identified RA-associated genes including *MED1, IL6R, CEP57, CDK6, RAD51B, RUNX1, RTKN2, RAG1, FADS1, RASGRP1, ETS1* and *COG6. miR-4308* showed seven candidate genes including *RTKN2, TRAF6, RAD51B, PTPN2, PLD4, TNFRSF14* and *SYNGR1. miR-3135b* showed 4 target genes including *GRHL2, CD28, PPIL4* and *RASGRP1.* Although a hypergeometric test showed miRNA targets significantly enriched in GWAS-identified RA candidate genes (P= 7.66×10^−3^, FC=1.59, **Figure 3B, Table S4**), permutation based analysis showed a non-significant enrichment (empirical P=0.23) indicating an unstable enrichment (**Figure S1B**). Finally, we also observed miRNA targets also enriched in FDA approved drug targets (P<2.2×10^−16^, empirical P=0.014, FC=1.82) indicating miRNA as the key regulatory network hubs might be promising drug targets for autoimmune diseases (**Figure 3C, Figure S1C**).

**Figure 3.**
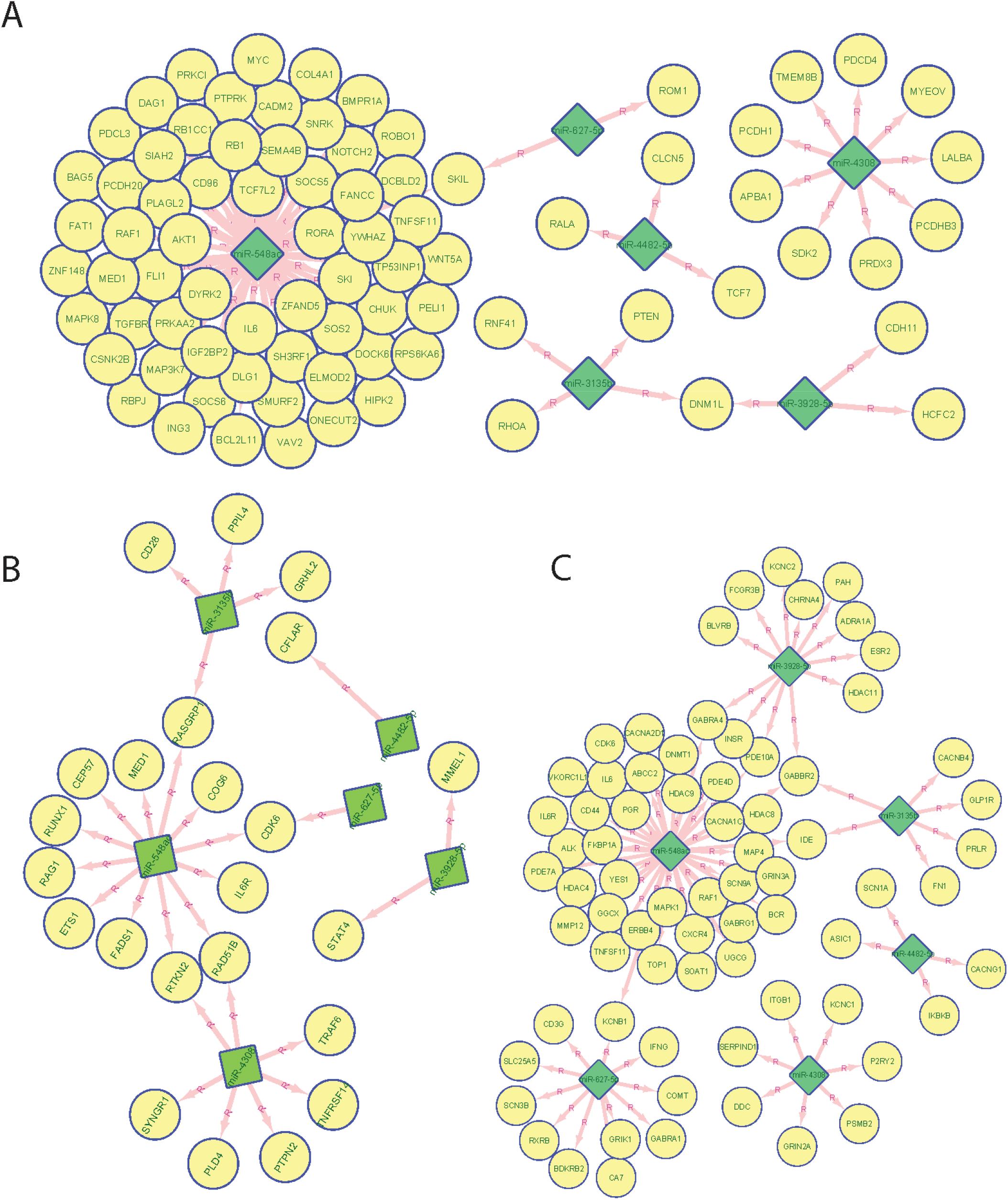
miRNA target analysis and target distribution in GWAS significant genes, immune genes and FDA drug target genes. (A) Network connection between the top 6 miRNA predicted regulatory targets and previous GWAS identified 123 RA associated genes. (B) Network connection between the top 6 miRNA predicted regulatory targets and 4,723 InnateDB collected immune genes. (C) Network connection between the top 6 miRNA predicted regulatory targets and 672 FDA approved drug target genes. miRNA was labelled as diamond filled with green color and gene were as circle filled with yellow color.

## Discussion

In this study, 225 East-Asian common SNPs located in human microRNA seed regions were genotyped and association and epistasis analyses were conducted to investigate the association between miRNA and seropositive RA in the Han Chinese population (Shanghai, China). To evaluate the potential association of these East-Asian-specific common miRNA SNPs with RA susceptibility, a case/control study was conducted involving 1,607 seropositive RA patients compared to 1,580 matched controls. The study identified 6 novel RA-associated miRNA SNPs (rs1414273 in *miR-548ac,* rs2620381 in *miR-627,* rs4285314 in *miR-3135b,* rs28477407 in *miR-4308;* rs5997893 in *miR-3928* and rs45596840 in *miR-4482*) and revealed the interaction between HLA alleles and miRNA SNPs, thereby advancing our understanding of RA pathogenesis.

We found rs45596840 and rs2620381 located in seed regions of *miR-4482* and *miR-627,* respectively, are significantly associated with RA. rs9268839-G (*HLA-DRB9*) was showed as a risk allele with OR=2.47 in Caucasians population and we found the OR is 1.72 indicating the consistent risk effect and might be required to be validated in African population. According to the miRNA-mRNA binding imputation, rs45596840 (*miR-4482*) and rs2620381 (*miR-627*) will affect more than 5,894 (2,132 gain and 3,517 loss, see *URL*) and 4,845 (2,593 gain and 2,252 loss, see *URL*) mRNA-miRNA bindings respectively. These miRNA target change might change the immune response significantly. In addition, the common regulation network between *miR-4707* and *miR-627* with *CD244, CAMTA1* might be an interesting. rs1414273 is located within the *miR-548ac* stem-loop sequencing in the first intron of the *CD58* gene, which is showed a strong linkage disequilibrium with the Multiple Sclerosis (MS)-associated haplotype.^46^ Hecker M et al. found that SNP rs1414273 might alter Drosha cleavage activity to provoke partial uncoupling of *CD58* gene expression and microRNA-548ac production from the shared primary transcript in immune cells and to regulate the inflammatory processes and the balance of protein folding and degradation.^47^ Although *CD58-CD2* interactions recruiting lymphocyte to inflammatory sites to play a crucial role in autoimmune disease.^48^ ^49^ Finally, in our previous research, we found DNA methylation of CD4+ cell in rheumatoid arthritis showed abnormal DNA methylation in HLA regions.^50^ In this current study, we found interactions between HLA-miRNA as contributing to RA susceptibility. Overall, the interaction between regulation/epigenetics and genetic variation in RA is a promising field for future research.

There are limitations with our study. Due to the scope of the study, only a limited number of ancestry-informative SNPs were used to control for confounding by population stratification. To validate the RA phenotyping, we utilized 4 East-Asian GWAS RA-associated variants to serve as positive controls for RA samples. We also required all the enrolled individuals were from three-generation Han Chinese family without genetic admixture with other non-Han ethnic individuals to avoid population stratification. This study did not provide functional validation of these miRNAs which is important to show biological validation of the miRNA findings and understand the mechanisms by which these miRNAs are involved in RA susceptibility. Subsequent studies should focus on functional studies of these miRNAs. Overall, we identified 6 miRNA SNPs which are significant associated with the Han Chinese RA population. Two of these SNPs are located in seed region of their miRNAs and are expected tol cause target gain/loss, while another three SNPs are located in the loop/pre-miRNA regions which may influence the stability of corresponding miRNAs.

## Supporting information

We found target genes of RA-associated miRNAs were significant enriched in the immune related gene category

Furthermore, we did not identify significant age, BMI, drinking and smoking history differences between cases and controls

We also conducted mode of inheritance-based association analyses (Table 2) and Cochran-Armitage Trend test-based association analyses

We found 19 SNP-SNP epistatic interactions with P<7.3x10-4, indicating significant interactions

Although a hypergeometric test showed miRNA targets significantly enriched in GWAS-identified RA candidate genes

## Declarations

### Acknowledgements

We thank all participating subjects for their kind cooperation in this study. All methods were performed in accordance with the relevant guidelines and regulations recorded in IRB approved research proposal. All patients were random enrolled and fulfilled the 2010 American College of Rheumatology classification criteria for RA.

### Contributors

SG and DH contributed to the conception, design and final approval of the submitted version. YJ, YB, JZ, QZ, TJ, YB, RZ, LX, CC, JS, XZ, YS, YQ, JC, XT, PC, QD, YZ, JL, QC, MG, ZL, WQ, YQ, YS, YS, HN developed consent form, recruited individuals, collected blood samples and clinical information collection. SG, YJ contributed to statistical data analysis. JS and DH contributed to the conception, study design, manuscript review and result explanation. The final manuscript was finally completed by SG, YJ, SS and DH. All authors read and approved the final manuscript.

### Patient and public involvement

Patients and/or the public were not involved in the design, or conduct, or reporting or dissemination plans of this research.

### Patient consent for publication

Not required.

### Ethical approval

On 15 April 2014, the study was reviewed and approved by Guanghua Hospital Research Institute Institutional Review Board (2015-SRA-01, Title: Genetic and Epigenetic Research to Rheumatoid Arthritis).

### Data availability statement

Summary information are access public and provided in supplementary table for further meta-analysis. In addition, the summary statistic has been deposited in the GitHub page https://github.com/Shicheng-Guo/miRNA-RA. Raw data and materials are available upon the request to the corresponding authors on reasonable collaboration request.

### URL

dbSNP153: https://ftp.ncbi.nlm.nih.gov/snp/latest_release/VCF/

miRBase: ftp://mirbase.org/pub/mirbase/CURRENT/README

miRNASNP: http://bioinfo.life.hust.edu.cn/miRNASNP2/index.php

miRDB: http://www.mirdb.org/ontology.html

InnateDB, 4,723 immune genes: https://www.innatedb.com/

rs2273626-target: http://bioinfo.life.hust.edu.cn/miRNASNP#!/snp?snp_id=rs2273626&location=Seed&one=1

rs45596840-target:http://bioinfo.life.hust.edu.cn/miRNASNP#!/snp?snp_id=rs45596840&location=Seed&one=1

rs2620381-targets: http://bioinfo.life.hust.edu.cn/miRNASNP#!/snp?snp_id=rs2620381&location=Seed&one=1

## Supplementary Matreial

**Figure S1. Re-sampling technique based enrichment analysis to miRNA regulatory targets.** *A random sampling technique was applied to assess the null distribution of p-values (n=50,000 iterations).37,875 total genes in the genome (GENCODE v32) were assumed for the calculations. Enrichment was measured with fold-change (FC) and p-values were calculated by summing the frequency of more extreme values in the null distribution of p-values.

**Table S1. Demographic information for the GHRA cohort**

**Table S2. Cochran-Armitage Trend test-based association analyses.**

**Table S3. Epistasis analysis to identify SNP-SNP interaction in rheumatoid arthritis susceptibility**

**Table S4. miRDB based miRNA target prediction for the 6 significant RA-miRNA.**

