## Supplementary material for "MicroRNA variants and HLA-miRNA interactions are novel rheumatoid arthritis susceptibility factors": We found target genes of RA-associated miRNAs were significant enriched in the immune related gene category

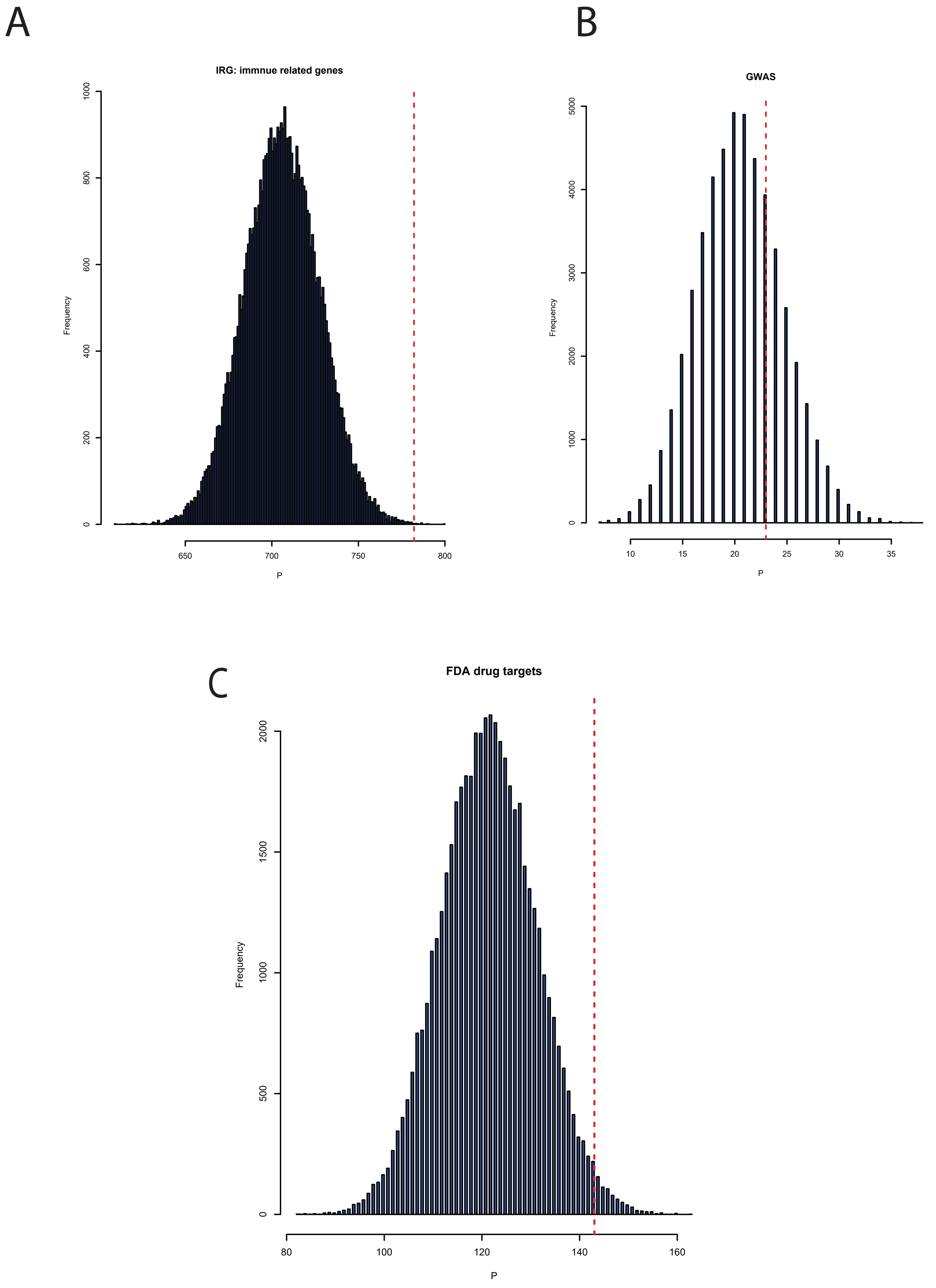


**Figure S1.Re-sampling technique based enrichment analysis to miRNA regulatory targets.** *A random sampling technique was applied to assess the null distribution of p-values (n=50,000 iterations).37,875 total genes in the genome (GENCODE v32) were assumed for the calculations. Enrichment was measured with fold-change (FC) and p-values were calculated by summing the frequency of more extreme values in the null distribution of p-values.
