## Supplementary material for "MicroRNA variants and HLA-miRNA interactions are novel rheumatoid arthritis susceptibility factors": Furthermore, we did not identify significant age, BMI, drinking and smoking history differences between cases and controls

Table S1. Demographic information for the GHRA cohort

| Characteristics | Case (n=1625) | Control (n=1598) | |
|---|---|---|---|
| | n | n | P-value |
| Age | | | |
| <60 | 786 | 721 | |
| ≥60 | 816 | 871 | P=0.035 |
| Sex | | | |
| Male | 318 | 314 | |
| Female | 1284 | 1278 | P=0.963 |
| BMI | | | |
| Underweight (BMI＜18.5) | 153 | 150 | |
| Normal(18.5≤BMI＜24) | 975 | 897 | |
| Overweight(24≤BMI＜28) | 387 | 433 | |
| Obesity (BMI≥28) | 84 | 112 | P=0.019 |
| Smoking | | | |
| Yes | 115 | 109 | |
| No | 1439 | 1483 | P=0.593 |
| Drinking | | | |
| Yes | 39 | 55 | |
| No | 1496 | 1537 | P=0.164 |

Chi-square test was applied in the differential distribution test and multiple test corrections were applied Bonferroni correction, therefore, 0.01 was used as the criterion for the significant differential distribution. Missing values for above characteristics were not showed in the table.
