## Supplementary material for "MicroRNA variants and HLA-miRNA interactions are novel rheumatoid arthritis susceptibility factors": We also conducted mode of inheritance-based association analyses (Table 2) and Cochran-Armitage Trend test-based association analyses

Table S2. Cochran-Armitage Trend test-based association analyses.

| CHR | SNP | A1 | A2 | AFF | UNAFF | P |
|---|---|---|---|---|---|---|
| 6 | rs9268839 | A | G | 1364/1850 | 1768/1392 | 1.439E-27 |
| 6 | rs9275376 | T | G | 804/2410 | 496/2664 | 1.804E-20 |
| 6 | rs9275375 | A | G | 1064/2150 | 788/2372 | 1.869E-12 |
| 6 | rs7752903 | G | T | 205/3009 | 136/3024 | 0.0002118 |
| 6 | rs4947332 | T | C | 72/3142 | 34/3126 | 0.0003235 |
| 1 | rs1414273 | C | T | 1303/1911 | 1412/1748 | 0.000911 |
| 15 | rs2620381 | C | A | 254/2960 | 318/2842 | 0.002716 |
| 14 | rs75330474 | T | C | 121/3093 | 159/3001 | 0.01286 |
| 18 | rs370878033 | G | A | 54/3160 | 81/3079 | 0.01553 |
| 7 | rs3823658 | A | G | 432/2782 | 363/2797 | 0.017 |
| 1 | rs74085143 | A | G | 130/3084 | 166/2994 | 0.02149 |
| 6 | rs7740161 | T | A | 459/2755 | 515/2645 | 0.02487 |
| 3 | rs4687672 | A | G | 1082/2132 | 983/2177 | 0.0292 |
| 22 | rs5997893 | A | G | 1509/1705 | 1565/1595 | 0.03957 |
| 19 | rs2967897 | T | C | 541/2673 | 473/2687 | 0.03959 |
| 15 | rs76468441 | T | C | 104/3110 | 76/3084 | 0.04463 |
| 1 | rs12402181 | A | G | 1157/2057 | 1064/2096 | 0.05407 |
| 19 | rs4112253 | C | G | 78/3136 | 102/3058 | 0.05698 |
| 14 | rs28477407 | T | C | 1290/1924 | 1341/1819 | 0.06512 |
| 11 | rs7350542 | G | A | 640/2574 | 574/2586 | 0.07059 |
| 11 | rs11237828 | C | T | 1139/2075 | 1052/2108 | 0.07137 |
| 19 | rs72996752 | G | A | 890/2324 | 814/2346 | 0.08222 |
| 3 | rs6771809 | C | T | 431/2783 | 378/2782 | 0.08625 |
| 3 | rs6787734 | C | T | 1110/2104 | 1156/2004 | 0.08912 |
| 16 | rs35613341 | C | G | 1426/1788 | 1336/1824 | 0.09094 |
| 10 | rs45596840 | A | G | 536/2678 | 577/2583 | 0.09535 |
| 19 | rs10406069 | A | G | 645/2569 | 687/2473 | 0.09964 |
| 7 | rs117344178 | T | C | 78/3136 | 98/3062 | 0.1035 |
| 16 | rs56292801 | G | A | 1427/1787 | 1341/1819 | 0.1119 |
| 15 | rs2060455 | A | G | 906/2308 | 836/2324 | 0.1167 |
| 8 | rs12549434 | C | T | 55/3159 | 71/3089 | 0.1238 |
| 6 | rs4285314 | G | A | 470/2744 | 506/2654 | 0.1287 |
| 11 | rs12803915 | A | G | 458/2756 | 493/2667 | 0.1311 |
| 17 | rs58390814 | T | C | 138/3076 | 112/3048 | 0.1312 |
| 12 | rs3751304 | C | T | 617/2597 | 654/2506 | 0.1328 |
| 2 | rs6717413 | G | A | 407/2807 | 362/2798 | 0.1334 |
| 5 | rs77966622 | A | G | 90/3124 | 70/3090 | 0.1354 |
| 3 | rs78790512 | A | G | 53/3161 | 68/3092 | 0.1408 |
| 3 | rs75715827 | C | T | 61/3153 | 77/3083 | 0.141 |
| 11 | rs174561 | C | T | 1317/1897 | 1238/1922 | 0.1426 |
| 12 | rs10878362 | A | C | 272/2942 | 236/2924 | 0.1459 |

| | | | | | | |
|---|---|---|---|---|---|---|
| 2 | rs74322818 | T | C | 73/3141 | 90/3070 | 0.147 |
| 4 | rs28664200 | C | T | 1595/1619 | 1512/1648 | 0.1484 |
| 19 | rs57111412 | G | A | 978/2236 | 910/2250 | 0.1491 |
| 1 | rs1953090 | G | T | 592/2622 | 538/2622 | 0.1515 |
| 14 | rs2273626 | C | A | 592/2622 | 700/2460 | 0.1618 |
| 19 | rs1003723 | T | C | 471/2743 | 503/2657 | 0.167 |
| 13 | rs1572687 | T | C | 923/2291 | 956/2204 | 0.1677 |
| 13 | rs67976778 | T | C | 1386/1828 | 1415/1745 | 0.1801 |
| 6 | rs67182313 | A | G | 801/2413 | 743/2417 | 0.1864 |
| 3 | rs2292181 | C | G | 111/3103 | 129/3031 | 0.194 |
| 1 | rs877722 | T | A | 641/2573 | 591/2569 | 0.2034 |
| 11 | rs174583 | T | C | 1340/1874 | 1268/1892 | 0.204 |
| 1 | rs73108496 | G | A | 329/2885 | 354/2806 | 0.2173 |
| 9 | rs3780662 | T | C | 270/2944 | 293/2867 | 0.2195 |
| 14 | rs145135208 | G | A | 268/2946 | 291/2869 | 0.2225 |
| 14 | rs12894467 | C | T | 787/2427 | 733/2427 | 0.2226 |
| 6 | rs17881225 | C | G | 915/2299 | 733/2427 | 0.2422 |
| 11 | rs515924 | G | A | 1097/2117 | 1122/2038 | 0.2438 |
| 5 | rs367805 | T | C | 1337/1877 | 1270/1890 | 0.2494 |
| 10 | rs641071 | T | G | 858/2356 | 884/2276 | 0.2501 |
| 1 | rs2275874 | T | C | 1139/2075 | 1163/1997 | 0.2554 |
| 16 | rs58353328 | G | A | 333/2881 | 301/2859 | 0.2629 |
| 17 | rs8078913 | C | T | 1056/2158 | 1080/2080 | 0.2687 |
| 21 | rs451887 | T | C | 414/2800 | 436/2724 | 0.2782 |
| 5 | rs11745587 | A | G | 817/2397 | 840/2320 | 0.2885 |
| 5 | rs702742 | G | A | 278/2936 | 297/2863 | 0.293 |
| 19 | rs8667 | A | G | 1467/1747 | 1402/1758 | 0.2966 |
| 7 | rs361399 | A | G | 598/2616 | 557/2603 | 0.3072 |
| 7 | rs361398 | A | G | 598/2616 | 557/2603 | 0.3072 |
| 7 | rs361397 | A | G | 598/2616 | 557/2603 | 0.3072 |
| 12 | rs11176006 | A | G | 319/2895 | 290/2870 | 0.3105 |
| 7 | rs62442513 | T | C | 261/2953 | 235/2925 | 0.312 |
| 2 | rs10175383 | C | G | 806/2408 | 759/2401 | 0.3215 |
| 12 | rs1683709 | G | A | 1559/1655 | 1494/1666 | 0.3327 |
| 17 | rs7208391 | G | C | 1500/1714 | 1437/1723 | 0.3419 |
| 19 | rs975947 | T | C | 1204/2010 | 1148/2012 | 0.3504 |
| 10 | rs7070684 | A | G | 1227/1987 | 1171/1989 | 0.3561 |
| 7 | rs11983381 | G | A | 284/2930 | 300/2860 | 0.3678 |
| 2 | rs62182086 | G | A | 129/3085 | 141/3019 | 0.3692 |
| 16 | rs16958290 | C | G | 1011/2203 | 1027/2133 | 0.3699 |
| 12 | rs1290910 | C | G | 1571/1643 | 1580/1580 | 0.37 |
| 12 | rs2289030 | C | G | 690/2524 | 707/2453 | 0.3804 |
| 12 | rs61938575 | A | G | 700/2514 | 717/2443 | 0.3842 |
| 12 | rs832733 | T | C | 650/2564 | 667/2493 | 0.3913 |

| | | | | | | |
|---|---|---|---|---|---|---|
| 4 | rs34115976 | G | C | 187/3027 | 200/2960 | 0.3961 |
| 12 | rs2682818 | A | C | 874/2340 | 831/2329 | 0.4153 |
| 19 | rs1688017 | A | G | 766/2448 | 726/2434 | 0.4156 |
| 5 | rs3734050 | T | C | 106/3108 | 116/3044 | 0.4173 |
| 11 | rs11032942 | C | T | 249/2965 | 262/2898 | 0.4206 |
| 6 | rs2276448 | C | T | 923/2291 | 879/2281 | 0.4267 |
| 20 | rs744591 | C | A | 1562/1652 | 1504/1656 | 0.4271 |
| 14 | rs61992671 | G | A | 91/3123 | 100/3060 | 0.4307 |
| 2 | rs2241347 | C | T | 730/2484 | 744/2416 | 0.4324 |
| 2 | rs2292879 | G | A | 1016/2198 | 1028/2132 | 0.4332 |
| 3 | rs78831152 | T | C | 333/2881 | 309/2851 | 0.4396 |
| 2 | rs58450758 | T | C | 559/2655 | 573/2587 | 0.4411 |
| 16 | rs2925980 | G | A | 1369/1845 | 1316/1844 | 0.4424 |
| 1 | rs79639536 | G | A | 134/3080 | 144/3016 | 0.4485 |
| 1 | rs619608 | A | G | 538/2676 | 551/2609 | 0.4539 |
| 10 | rs7911488 | G | A | 1085/2129 | 1039/2121 | 0.4581 |
| 10 | rs2368393 | G | A | 1049/2165 | 1005/2155 | 0.4753 |
| 10 | rs2368392 | A | G | 1049/2165 | 1005/2155 | 0.4753 |
| 2 | rs79402775 | A | G | 440/2774 | 452/2708 | 0.4827 |
| 5 | rs266435 | C | G | 521/2693 | 492/2668 | 0.4875 |
| 7 | rs6464546 | A | G | 476/2738 | 449/2711 | 0.4993 |
| 1 | rs76756293 | T | C | 314/2900 | 324/2836 | 0.5159 |
| 1 | rs17111728 | C | T | 316/2898 | 326/2834 | 0.5166 |
| 3 | rs142342924 | A | G | 750/2464 | 716/2444 | 0.5212 |
| 14 | rs58834075 | T | C | 81/3133 | 72/3088 | 0.5232 |
| 5 | rs62376934 | A | G | 325/2889 | 335/2825 | 0.529 |
| 4 | rs73239138 | A | G | 1259/1955 | 1262/1898 | 0.5292 |
| 2 | rs77373668 | C | G | 242/2972 | 251/2909 | 0.5339 |
| 5 | rs936581 | A | G | 54/3160 | 47/3113 | 0.5344 |
| 8 | rs66683138 | A | G | 1293/1921 | 1247/1913 | 0.5371 |
| 3 | rs10934682 | G | T | 536/2678 | 545/2615 | 0.5401 |
| 19 | rs56061231 | G | A | 1474/1740 | 1425/1735 | 0.5439 |
| 10 | rs12416605 | T | C | 105/3109 | 95/3065 | 0.5462 |
| 3 | rs9842591 | A | C | 1606/1608 | 1555/1605 | 0.5468 |
| 6 | rs77651740 | T | G | 1157/2057 | 1115/2045 | 0.5472 |
| 14 | rs2296319 | A | G | 419/2795 | 428/2732 | 0.5531 |
| 1 | rs701213 | C | T | 1029/2185 | 1032/2128 | 0.5842 |
| 9 | rs56195815 | T | C | 345/2869 | 326/2834 | 0.5897 |
| 19 | rs7247767 | G | A | 1289/1925 | 1288/1872 | 0.5928 |
| 2 | rs56148568 | C | T | 670/2544 | 642/2518 | 0.5995 |
| 7 | rs60871950 | A | G | 761/2453 | 731/2429 | 0.6026 |
| 19 | rs75598818 | A | G | 334/2880 | 341/2819 | 0.6042 |
| 8 | rs78360334 | C | T | 278/2936 | 262/2898 | 0.6052 |
| 8 | rs56863230 | C | G | 221/2993 | 207/2953 | 0.606 |

| | | | | | | |
|---|---|---|---|---|---|---|
| 19 | rs7247237 | T | C | 1289/1925 | 1287/1873 | 0.6106 |
| 8 | rs10505168 | C | T | 1481/1733 | 1476/1684 | 0.6156 |
| 17 | rs12451747 | A | C | 1520/1694 | 1475/1685 | 0.6184 |
| 11 | rs2155248 | G | T | 192/3022 | 198/2962 | 0.6211 |
| 1 | rs45530340 | T | C | 458/2756 | 464/2696 | 0.6212 |
| 11 | rs2986407 | T | C | 598/2616 | 603/2557 | 0.6254 |
| 8 | rs404337 | G | A | 1484/1730 | 1478/1682 | 0.6265 |
| 8 | rs75404472 | T | C | 265/2949 | 250/2910 | 0.6294 |
| 17 | rs111756476 | T | A | 370/2844 | 376/2784 | 0.6298 |
| 8 | rs6997249 | A | G | 354/2860 | 360/2800 | 0.6307 |
| 1 | rs75036690 | A | G | 101/3113 | 106/3054 | 0.6326 |
| 8 | rs28655823 | C | G | 532/2682 | 537/2623 | 0.6345 |
| 2 | rs10192411 | G | A | 240/2974 | 246/2914 | 0.6371 |
| 2 | rs6726779 | C | T | 856/2358 | 858/2302 | 0.639 |
| 2 | rs6430498 | A | G | 1141/2073 | 1139/2021 | 0.6483 |
| 10 | rs72810954 | A | G | 61/3153 | 65/3095 | 0.6504 |
| 16 | rs74469188 | C | T | 229/2985 | 216/2944 | 0.6535 |
| 8 | rs487571 | T | C | 1448/1766 | 1406/1754 | 0.6539 |
| 6 | rs12197631 | G | T | 152/3062 | 157/3003 | 0.6552 |
| 11 | rs12801172 | C | G | 1488/1726 | 1480/1680 | 0.6676 |
| 15 | rs7162033 | G | C | 595/2619 | 572/2588 | 0.6688 |
| 15 | rs7183051 | A | G | 595/2619 | 572/2588 | 0.6688 |
| 5 | rs2042253 | C | T | 1346/1868 | 1307/1853 | 0.6749 |
| 9 | rs35196866 | C | A | 778/2436 | 751/2409 | 0.6773 |
| 1 | rs701214 | T | C | 342/2872 | 346/2814 | 0.6876 |
| 16 | rs57629257 | T | C | 1335/1879 | 1327/1833 | 0.7143 |
| 22 | rs4822739 | G | C | 723/2491 | 723/2437 | 0.7146 |
| 4 | rs12512664 | G | A | 110/3104 | 103/3057 | 0.7165 |
| 6 | rs7769202 | C | T | 705/2509 | 705/2455 | 0.7178 |
| 17 | rs6505162 | A | C | 582/2632 | 583/2577 | 0.7248 |
| 4 | rs1077020 | C | T | 854/2360 | 852/2308 | 0.7256 |
| 5 | rs62376935 | T | C | 921/2293 | 893/2267 | 0.727 |
| 22 | rs4078443 | T | C | 735/2479 | 711/2449 | 0.7276 |
| 9 | rs58605477 | G | A | 406/2808 | 408/2752 | 0.7356 |
| 4 | rs77639117 | T | A | 146/3068 | 149/3011 | 0.7458 |
| 11 | rs67042258 | A | G | 494/2720 | 495/2665 | 0.7465 |
| 20 | rs6513497 | G | T | 338/2876 | 340/2820 | 0.7506 |
| 17 | rs745666 | C | G | 1126/2088 | 1119/2041 | 0.7521 |
| 12 | rs3817551 | G | T | 1357/1857 | 1322/1838 | 0.7567 |
| 16 | rs12708966 | A | G | 82/3132 | 77/3083 | 0.7663 |
| 10 | rs12780876 | A | T | 571/2643 | 570/2590 | 0.7749 |
| 17 | rs73410309 | C | G | 241/2973 | 243/2917 | 0.7757 |
| 19 | rs10423365 | G | A | 942/2272 | 936/2224 | 0.7875 |
| 2 | rs243080 | G | A | 1073/2141 | 1065/2095 | 0.7919 |

| | | | | | | |
|---|---|---|---|---|---|---|
| 2 | rs2292832 | C | T | 880/2334 | 856/2304 | 0.7919 |
| 5 | rs10061133 | G | A | 828/2386 | 805/2355 | 0.793 |
| 19 | rs895819 | C | T | 835/2379 | 830/2330 | 0.7975 |
| 8 | rs2114358 | G | A | 802/2412 | 780/2380 | 0.8018 |
| 5 | rs12523324 | G | A | 1338/1876 | 1325/1835 | 0.8074 |
| 7 | rs4909237 | T | C | 754/2460 | 749/2411 | 0.8182 |
| 12 | rs11614913 | C | T | 1485/1729 | 1451/1709 | 0.8184 |
| 5 | rs3112399 | A | T | 415/2799 | 402/2758 | 0.8215 |
| 10 | rs11014002 | T | C | 672/2542 | 654/2506 | 0.8314 |
| 2 | rs71428439 | G | A | 545/2669 | 542/2618 | 0.8351 |
| 15 | rs1439619 | G | T | 929/2285 | 906/2254 | 0.8356 |
| 16 | rs7500280 | T | C | 1249/1965 | 1220/1940 | 0.8362 |
| 1 | rs72646786 | T | C | 506/2708 | 492/2668 | 0.8466 |
| 11 | rs116971067 | C | G | 128/3086 | 123/3037 | 0.8526 |
| 1 | rs2070960 | T | C | 614/2600 | 598/2562 | 0.8548 |
| 16 | rs7499278 | G | A | 1245/1969 | 1217/1943 | 0.855 |
| 11 | rs670637 | G | A | 62/3152 | 59/3101 | 0.856 |
| 11 | rs634171 | T | A | 62/3152 | 59/3101 | 0.856 |
| 15 | rs4414449 | G | A | 1115/2099 | 1103/2057 | 0.8578 |
| 15 | rs4577031 | A | T | 1115/2099 | 1103/2057 | 0.8578 |
| 17 | rs7210937 | C | G | 1391/1823 | 1361/1799 | 0.8642 |
| 19 | rs10412196 | C | T | 261/2953 | 253/2907 | 0.8674 |
| 22 | rs60308683 | A | G | 142/3072 | 137/3023 | 0.8712 |
| 17 | rs7207008 | A | T | 924/2290 | 903/2257 | 0.8796 |
| 17 | rs620301 | G | A | 1507/1707 | 1476/1684 | 0.8866 |
| 6 | rs68035463 | A | C | 464/2750 | 460/2700 | 0.8915 |
| 6 | rs67106263 | A | G | 464/2750 | 460/2700 | 0.8915 |
| 10 | rs2043556 | C | T | 851/2363 | 832/2328 | 0.8943 |
| 5 | rs257095 | C | T | 115/3099 | 115/3045 | 0.8944 |
| 10 | rs4919510 | C | G | 1377/1837 | 1349/1811 | 0.9008 |
| 16 | rs77250474 | A | G | 87/3127 | 84/3076 | 0.9047 |
| 12 | rs12314280 | C | T | 340/2874 | 337/2823 | 0.9107 |
| 5 | rs7709117 | G | A | 1561/1653 | 1539/1621 | 0.914 |
| 16 | rs897984 | T | C | 228/2986 | 222/2938 | 0.9144 |
| 20 | rs6513496 | C | T | 387/2827 | 378/2782 | 0.9226 |
| 15 | rs2168518 | A | G | 498/2716 | 487/2673 | 0.9253 |
| 3 | rs1514422 | A | G | 885/2329 | 867/2293 | 0.9293 |
| 3 | rs35452137 | C | A | 151/3063 | 147/3013 | 0.9305 |
| 10 | rs7896283 | G | A | 1415/1799 | 1388/1772 | 0.9347 |
| 1 | rs521188 | G | A | 989/2225 | 975/2185 | 0.9426 |
| 19 | rs10422347 | T | C | 116/3098 | 113/3047 | 0.9431 |
| 18 | rs12456845 | C | T | 863/2351 | 851/2309 | 0.944 |
| 12 | rs17022749 | T | C | 445/2769 | 436/2724 | 0.9554 |
| 16 | rs74656628 | A | T | 129/3085 | 126/3034 | 0.9577 |

| | | | | | | |
|---|---|---|---|---|---|---|
| 20 | rs3746444 | G | A | 459/2755 | 450/2710 | 0.9631 |
| 11 | rs550894 | A | C | 803/2411 | 788/2372 | 0.9653 |
| 6 | rs9295535 | C | T | 992/2222 | 974/2186 | 0.9706 |
| 19 | rs3745198 | G | C | 1515/1699 | 1491/1669 | 0.9707 |
| 2 | rs56239160 | A | G | 1095/2119 | 1078/2082 | 0.9708 |
| 17 | rs9913045 | A | G | 785/2429 | 773/2387 | 0.9718 |
| 14 | rs2296320 | A | G | 153/3061 | 151/3009 | 0.9731 |
| 17 | rs66507245 | A | T | 1567/1647 | 1542/1618 | 0.9732 |
| 3 | rs9877402 | G | A | 150/3064 | 148/3012 | 0.9753 |
| 12 | rs75258105 | T | G | 338/2876 | 333/2827 | 0.9775 |
| 7 | rs7804972 | A | G | 466/2748 | 458/2702 | 0.995 |
| 9 | rs13299349 | A | G | 348/2866 | 342/2818 | 0.995 |

* hg19 genome coordinate
