## Supplementary material for "MicroRNA variants and HLA-miRNA interactions are novel rheumatoid arthritis susceptibility factors": We found 19 SNP-SNP epistatic interactions with P<7.3x10-4, indicating significant interactions

Table S3. Epistasis analysis to identify SNP-SNP interaction in rheumatoid arthritis susceptibility

| CHR1 | SNP1 | miRNA-1 | CHR2 | SNP2 | miRNA-2 | OR | P |
|---|---|---|---|---|---|---|---|
| 1 | rs1414273 | MIR548AC | 7 | rs117344178 | MIR595 | 0.313929 | 8.76E-06 |
| 8 | rs6997249 | MIR3686 | 14 | rs2273626 | MIR4707 | 0.544402 | 8.79E-06 |
| 8 | rs2114358 | MIR1206 | 15 | rs76468441 | MIR548AP | 0.274239 | 1.27E-05 |
| 6 | rs4285314 | MIR3135B | 16 | rs16958290 | MIR8058 | 1.53597 | 4.02E-05 |
| 5 | rs367805 | MIR3936 | 10 | rs641071 | MIR4482 | 0.716913 | 6.18E-05 |
| 7 | rs7804972 | MIR6839 | 17 | rs620301 | MIR548BC | 1.47573 | 0.000134 |
| 14 | rs2296320 | MIR4706 | 22 | rs5997893 | MIR3928 | 1.93891 | 0.000134 |
| 5 | rs2042253 | MIR5197 | 6 | rs9295535 | MIR5689HG | 1.34491 | 0.000174 |
| 10 | rs7070684 | MIR548AK | 11 | rs2155248 | MIR1304 | 1.83627 | 0.00026 |
| 1 | rs73108496 | MIR4428 | 11 | rs67042258 | MIR6128 | 0.558997 | 0.000376 |
| 2 | rs2241347 | MIR3130 | 5 | rs7709117 | MIR4634 | 1.35183 | 0.00044 |
| 1 | rs877722 | MIR4671 | 5 | rs62376935 | MIR585 | 0.700077 | 0.000473 |
| 8 | rs75404472 | MIR3674 | 16 | rs2925980 | MIR7854 | 0.634786 | 0.000485 |
| 9 | rs3780662 | MIR4672 | 10 | rs11014002 | MIR603 | 0.580794 | 0.000526 |
| 7 | rs4909237 | MIR595 | 11 | rs12801172 | MIR10392 | 1.3425 | 0.000583 |
| 2 | rs10175383 | MIR3679 | 8 | rs10505168 | MIR2053 | 0.748224 | 0.000612 |
| 9 | rs56195815 | MIR548AW | 10 | rs7911488 | MIR1037 | 1.53774 | 0.000639 |
| 3 | rs6771809 | MIR6826 | 14 | rs12894467 | MIR300 | 0.656563 | 0.000676 |
| 17 | rs58390814 | MIR6780A | 19 | rs56061231 | MIR6795 | 1.9326 | 0.000734 |
| 3 | rs75715827 | MIR944 | 16 | rs57629257 | MIR1972 | 0.428358 | 0.000847 |
| 6 | rs68035463 | MIR3144 | 14 | rs12894467 | MIR300 | 1.50915 | 0.000933 |

miRNA genomic position was obtained from miRBase 22 which is based on hg38 and then was liftover to hg19 for the further analysis. Genomic position in the table is hg19 version.
