## Supplementary material for "MicroRNA variants and HLA-miRNA interactions are novel rheumatoid arthritis susceptibility factors": Although a hypergeometric test showed miRNA targets significantly enriched in GWAS-identified RA candidate genes

Table S4. miRDB based miRNA target prediction for the 6 significant RA-miRNA.

| Target Detail | Target Rank | Target Score | miRNA Name | Gene Symbol |
|---|---|---|---|---|
| Details | 1 | 100 | hsa-miR-548ac | CREBBP |
| Details | 2 | 100 | hsa-miR-548ac | UBE2D1 |
| Details | 3 | 100 | hsa-miR-548ac | ZCCHC14 |
| Details | 4 | 100 | hsa-miR-548ac | RSBN1 |
| Details | 5 | 99 | hsa-miR-548ac | PSD3 |
| Details | 6 | 99 | hsa-miR-548ac | ARFGEF3 |
| Details | 7 | 99 | hsa-miR-548ac | MYC |
| Details | 8 | 99 | hsa-miR-548ac | GFPT1 |
| Details | 9 | 99 | hsa-miR-548ac | GRPEL2 |
| Details | 10 | 99 | hsa-miR-548ac | ONECUT2 |
| Details | 11 | 99 | hsa-miR-548ac | MAPK8 |
| Details | 12 | 99 | hsa-miR-548ac | DCBLD2 |
| Details | 13 | 99 | hsa-miR-548ac | IKZF2 |
| Details | 14 | 99 | hsa-miR-548ac | ITPRIPL2 |
| Details | 15 | 99 | hsa-miR-548ac | CCNYL1 |
| Details | 16 | 99 | hsa-miR-548ac | SCYL2 |
| Details | 17 | 99 | hsa-miR-548ac | PDS5A |
| Details | 18 | 99 | hsa-miR-548ac | INPP4A |
| Details | 19 | 99 | hsa-miR-548ac | DYRK2 |
| Details | 20 | 98 | hsa-miR-548ac | TCF7L2 |

Please check more targets:
https://github.com/Shicheng-Guo/miRNA-RA/blob/master/SupplementaryTable/Table-S4.xlsx
